# A Cryptic Prophage Transcription Factor Drives Phenotypic Changes via Host Gene Regulation

**DOI:** 10.1101/2024.09.21.614188

**Authors:** P. Lally, V.H. Tierrafría, L. Gómez-Romero, A. Stringer, J. Collado-Vides, J.T. Wade, J.E. Galagan

## Abstract

Cryptic prophages (CPs) are elements of bacterial genomes acquired from bacteriophage that infect the host cell and ultimately become stably integrated within the host genome. While some proteins encoded by CPs can modulate host phenotypes, the potential for Transcription Factors (TFs) encoded by CPs to impact host physiology by regulating host genes has not been thoroughly investigated. In this work, we report hundreds of host genes regulated by DicC, a DNA-binding TF encoded in the Qin prophage of *Esherichia coli*. We identified host-encoded regulatory targets of DicC that could be linked to known phenotypes of its induction. We also demonstrate that a DicC-induced growth defect is largely independent of other Qin prophage genes. Our data suggest a greater role for cryptic prophage TFs in controlling bacterial host gene expression than previously appreciated.

## Introduction

Prophages are lysogenic bacteriophage genomes that have integrated into a bacterial genome [1-3]. In the lysogenic state, the prophage remains inactive and replicates alongside the host genome, but in the lytic state the prophage creates bacteriophage progeny and eventually lyses the host cell. The balance between these states is coordinated by transcription factors (TFs) [4, 5], proteins that regulate transcription of genes via binding to DNA. Mutations in the prophage region or genome rearrangement can render the prophage incapable of activating the lytic state, causing it to become stably integrated into the host genome [2]. Although these “cryptic prophages” (CPs) are often considered dormant passengers inside their host, prior reports have demonstrated an impact of CPs on the host ability to respond to cellular stresses [6, 7]. For example, the 9 cryptic prophages in *Escherichia coli* have been shown to protect it from peroxide, antibiotic, and acid treatments [7]. However, the specific mechanisms by which these phenotypes are produced are not fully understood.

The Qin cryptic prophage has previously been shown to assist *E. coli*’s ability to resist stress associated with β-lactam antibiotics, oxidative stress, and acid stress [7]. A further study suggested that resistance is facilitated by pushing cells into a “Viable But Not Culturable” (VBNC) state, where cells are not actively growing but are nonetheless still alive [6]. Moreover, the same study also suggested that expression of the prophage’s *dicB* and *dicF* genes, which encode a small protein and small RNA, respectively, prevent entry into the VBNC state, and repress stress resistance. DicB inhibits septal ring formation by targeting FtsZ through an interaction with MinC, both encoded by host genes, which ceases cell division leading to death, while the *dicF* RNA inhibits translation of FtsZ [8, 9]. Expression of *dicB* and *dicF* is regulated by two prophage-encoded transcription factors. The Qin-encoded TF DicA directly represses transcription of both *dicB* and *dicF* [10]. An additional Qin-encoded TF, DicC, directly represses *dicA* [6]. DicC has also been suggested to induce expression of *dicB* and *dicF*, although it is not known if this regulation is direct, or indirect via repression of *dicA* [6]. A growth defect was also observed under *dicC* induction, and was suggested to be a consequence of *dicC* activating expression of *dicB* [6]. It has been suggested that *dicA* and *dicC* act as a bistable switch and master regulators of Qin prophage genes [10]; however, little is known about the broader transcriptional regulatory potential of *dicA* and *dicC*.

Here, we show that DicC has the potential for widespread transcriptional regulatory control of host-encoded genes. We show that DicC binds hundreds of regions across the *E. coli* genome, enriched for intergenic regions. We also show that DicC induction leads to differential expression of large numbers of genes, and we identify several directly regulated host genes that could drive the decrease in acid and oxidative stress resistances, as well as the growth defect seen with *dicC* induction. Finally, we show that *dicC* induction drives the growth defect independently of *dicB* and *dicF*, as well as the entire Qin prophage. Overall, our results suggest a role for prophage-encoded TFs to directly regulate a wide range of bacterial host genes to modulate host physiology.

## Results

### Inducible ChIP-Seq Reveals Widespread DicC Binding

The Qin prophage of *E. coli* encompasses 45 genes including *dicC* and *dicA*, which are transcribed divergently from the same intergenic region (Figure 1A). *dicC* is operonic with *ydfX*, which encodes a protein of unknown function, and *intK*, which increases resistance to macrolide antibiotics [11]. Downstream of *dicA* is a promoter capable of transcribing *ydfABC*, and potentially *dicF, dicB*, and *ydfD* [12]. There is also a promoter upstream of *dicF* that can transcribe it alone, and a promoter upstream of *dicB* capable of transcribing it along with *ydfD*, and pseudogenes *ydfE, insD7*, and *intQ* (not pictured) [12]. While functions for *ydfABC* are not known, *dicF* encodes a small RNA that inhibits cell division by preventing *ftsZ* translation [9], *dicB* encodes a protein that inhibits cell division by enhancing MinC’s inhibitory activity on FtsZ [8], and *ydfD* encodes a protein capable of lysing cells on induction, although lysis is prevented by co-expression of *dicB* [13]. DicA represses transcription of *dicC* and its operon, as well as the genes transcribed from the *dicB* promoter [10, 12]. DicA-mediated repressions of both *dicC* and *dicB* can be relieved by the Qin gene *rem* through an unknown mechanism [10]. Additionally, YjdC, a TF encoded by the *E. coli* host, regulates DicA targets in the absence of DicA [14].

**Figure 1.**
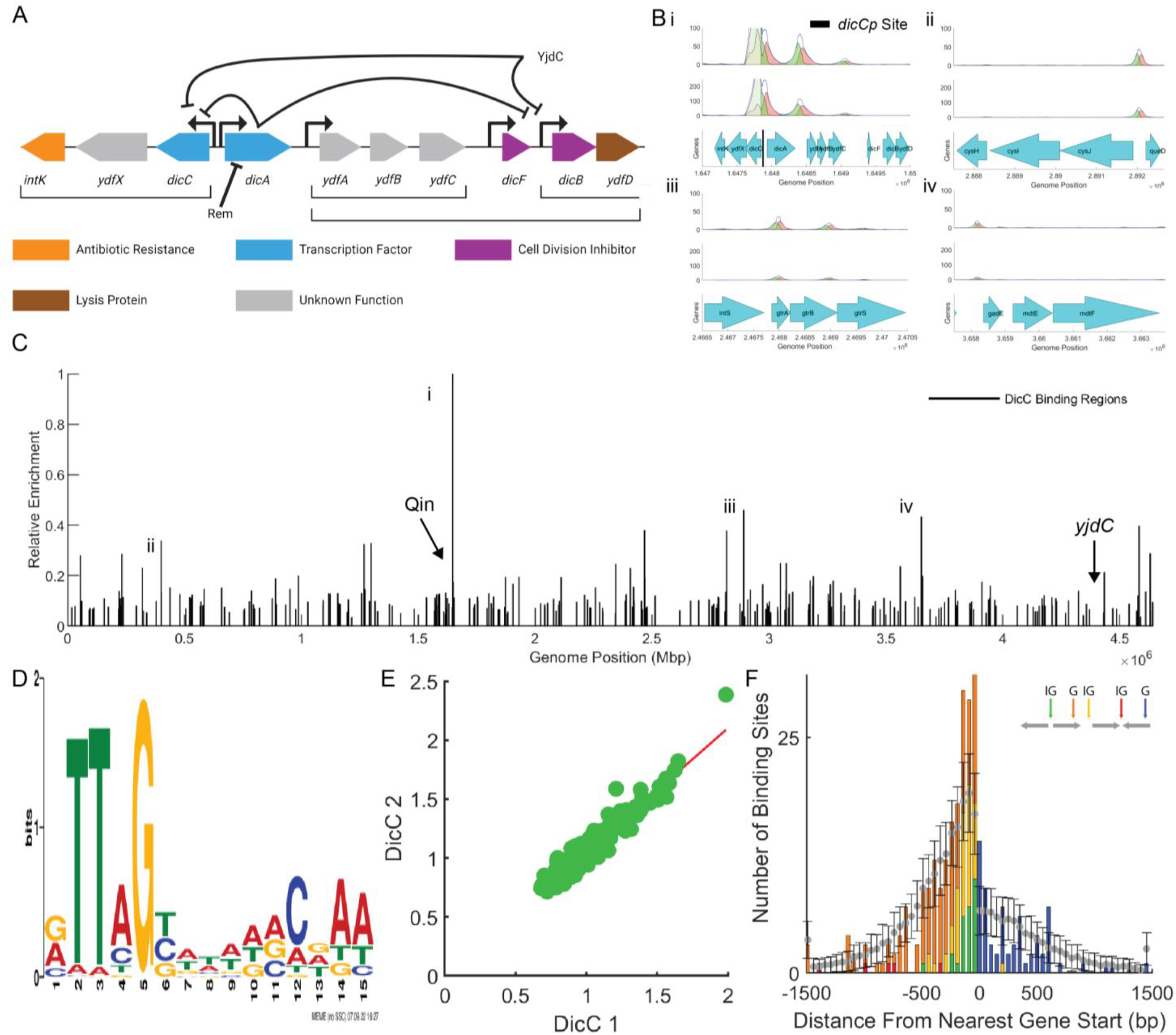
ChIP-Seq of Inducible DicC Reveals Widespread Genomic Binding. **A)** DicC is transcribed from a divergent promoter with *dicC* on one side and *dicA* on the other. DicA is known to repress *dicC* by binding to the divergent promoter, DicA is additionally known to repress the genes downstream of it including *ydfABC* and *dicFB*. An native (non-prophage) *E. coli* TF, YjdC, has been shown capable of repressing *dicA*’s targets in its absence as well. **B)** Coverage plots showing ChIP-Seq coverage of two DicC replicates at: **i)** overlapping the known *dicA* binding site (black bar) at the dicACp promoter region shown in (A); **ii)** upstream of *cysJIH*; **iii)** inside of another prophage (CPS-54); **iv)** upstream of another TF, *gadE*. **C)** Genome-wide DicC binding regions. Each bar represents a region identified from our ChIP-Seq, with its height indicating the relative enrichment of that region compared to the most highly-enriched region. **D)** MEME motif generated from DicC-bound regions shows DicC prefers palindromic sequences with TT(A/C/T)G on one side, and C(G/A/T)AA on the other. **E)** Enrichment of bound regions across replicates of DicC ChIP-Seq are highly correlated. **F)** Location of binding regions relative to start position of the nearest gene (G – genic, IG – intergenic). Binding regions between 150 bp upstream and 50 bp downstream are overrepresented relative to random expectation (grey points), regions > 1.5 kb up-or downstream of the nearest gene are grouped into a single bar on either edge.

In a separate study, we generated a compendium of TF binding profiles for *E. coli* TFs using ChIP-Seq. Through this study, we identified 279 bound regions for DicC using a strain that induces *dicC* expression from a plasmid [15]. The strongest binding for DicC occurs at its own promoter in the Qin prophage, where we infer two DicC binding sites, one on either side of *dicA* (Figure 1Bi). An additional weak binding region was observed upstream of *dicF*. The second-strongest binding region for DicC was found upstream of the *cysJIH* genes that function in sulfate assimilation (Figure 1Bii). We also observed DicC binding regions inside other *E. coli* prophages, including CPS-54 (Figure 1Biii), and upstream of the central activator of the glutamate-dependent acid resistance (GDAR) system, *gadE* (Figure 1Biv).

A genome-wide plot of each identified DicC-bound region with its enrichment relative to the most strongly bound region is shown in Figure 1C. The vast majority of identified DicC-bound regions displayed weaker binding, with 167/279 regions < 10% enriched relative to the most strongly bound region. Nonetheless, weak binding has been shown capable of driving significant phenotypic changes [16-18]. DicC-bound regions are strongly enriched for a palindromic sequence (Figure 1D). Enrichment of bound regions was highly reproducible across ChIP-Seq replicates (Figure 1E), and regions were strongly enriched for locations within 150 bp upstream and 50 bp downstream of gene starts (two-sample Kolmogorov-Smirnov p-value 0.0036; Figure 1F). Intergenic binding regardless of proximity to gene starts was also strongly enriched (34.78% of DicC regions vs 10.96% of genomic DNA, ∼3.4-fold overrepresented vs ∼2.5-fold for TFs on average). Intergenic regions often contain promoters, suggesting the potential for DicC to regulate many host genes.

### Evidence for Direct and Indirect Regulation of Host Gene Transcription by DicC

To test whether DicC regulates gene expression, we performed RNA-Seq to compare RNA levels in *E. coli* expressing *dicC* from an inducible plasmid relative to cells expressing an empty plasmid. Thus, we identified differentially expressed (DE) genes (|log_2_ fold-change| > 2, FDR < 0.05) at 2, 5, and 24 hours post-induction of DicC. We detected 31, 934, and 9 genes DE at 2, 5, and 24 hours post DicC induction, respectively, for a total of 939 DE genes across all time points. Only 4 of these genes are members of the Qin prophage (*dicA, ydfA, ydfJ*, and *ydfO*), demonstrating that DicC is capable of broad regulation of host-encoded genes.

We sought to identify instances of direct DicC regulation by associating DE genes with DicC-bound regions from our ChIP-Seq data. Thus, we assigned putative direct target genes to each DicC-bound region by identifying the nearest transcription unit (TU) on each side of a DicC-bound region and associating each gene within the corresponding TUs as a potential direct DicC target (Methods), providing a total of 765 potential targets. At 2, 5, and 24 hours we found 18 (2.4%), 195 (25.5%), and 2 (0.26%) of the 765 target genes were DE, respectively (Figure 2). Across all time points, 196 genes were both targets and DE (Supplementary Figure 1). Differentially expressed genes were enriched for binding targets at 2 and 5 hours, but not 24 hours (Fisher’s exact p-values 0.00049, 0.0049, and 0.66, respectively). In the cases of 2 and 5 hours, we would have expected roughly 5 and 165 direct DE genes if binding and DE were independent. These findings suggest that DicC could be capable of directly regulating nearly 200 host-encoded genes.

**Figure 2.**
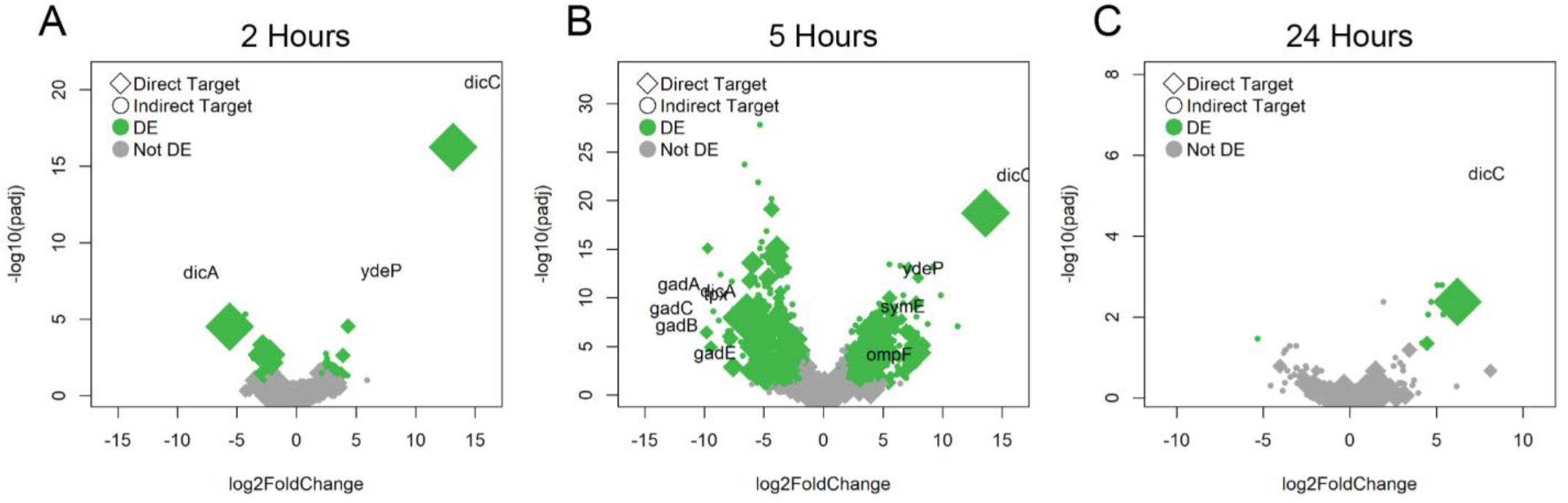
DicC Induction Drives Differential Expression of Genes. Volcano plots of DicC-induced expression at 2-(left), 5-(middle), and 24-hours (right). The x-axis represents log_2_ fold-change under DicC induction compared to induction of an empty plasmid. The y-axis shows the -log_10_(adjusted p-value) of each point. Diamonds represent genes with a nearby DicC binding site and the size of the point is relative to the strength of the site. Differentially expressed genes are plotted in green, and non-DE genes in gray.

While the enriched association of DicC binding sites with DE genes suggests some degree of direct regulation, indirect regulation could occur through regulation of other TFs. We found two DE TFs that could be associated with DicC binding sites at 2 hours, and 15 at 5 hours. One of these TFs was *dicA*, which was down-regulated 48-fold at 2 hours and 84-fold at 5 hours (but not DE at 24 hours). This is consistent with other findings that DicC represses *dicA* [6, 10]. Interestingly, differential expression of *dicF* and *dicB* was not detected at any time point, contrary to previous findings suggesting DicC regulates *dicF* and *dicB* expression [6, 19]. A table of fold changes and adjusted p-values for all genes is included in supplementary material.

To gain insight into potential roles of DicC-regulated genes, we performed a Gene Ontology (GO) Enrichment Analysis using PANTHER [20] (Supplementary Material). We identified statistically significant enrichment (FDR < 0.05) of GOs involved in intracellular pH elevation (Glutamate-Dependent Acid Resistance *gadABCE* genes; as well as *adiA, kefC*, and *kefF*, the former of which helps pH maintenance by degradation of arginine, and the latter two facilitate potassium:proton antiport). We also found enrichment of oxidative stress genes, including *tpx* and 35 others. While this analysis provides some evidence for selection of certain bioprocesses, the enriched ontologies spanned from pH regulation and stress resilience to biofilm formation and metabolic processes of various compounds (including trehalose, galactitol, and glyoxylate). Additionally, some GO enrichment could have been due to operons often containing functionally related genes, where a single regulatory event could result in enrichment of an entire category. Overall, the wide range of enriched ontologies suggests that DicC may regulate host genes broadly rather than specifically selecting genes of certain cellular pathways.

### Presence of Qin Toxins DicB, DicF, and YdfD Have Minimal Influence on DicC-Induced Expression Changes

To determine whether regulation of host-encoded genes via DicC induction could be a consequence of expression of the toxic *dicB, dicF*, and *ydfD* genes, we performed RNA-Seq comparing wild-type and Δ*dicBF*-*ydfD* strains with DicC induction to a wild-type strain with an induced empty plasmid. We saw statistically significant agreement in the direction of expression changes between wild-type and Δ*dicBF*-*ydfD* backgrounds at each time point (Figure 3; Chi-squared p-value < 2.2×10^−16^ at each time point post-induction). When we looked at the genes that were DE in the wild-type background at each time point, the direction of regulation was always consistent in the Δ*dicBF*-*ydfD* background. Overall, these data suggest that the differential expression of host-encoded genes observed with *dicC* induction was not dependent on the presence of the toxic *dicB, dicF*, and *ydfD* genes, the former two of which *dicC* has been suggested to activate [6, 19].

**Figure 3.**
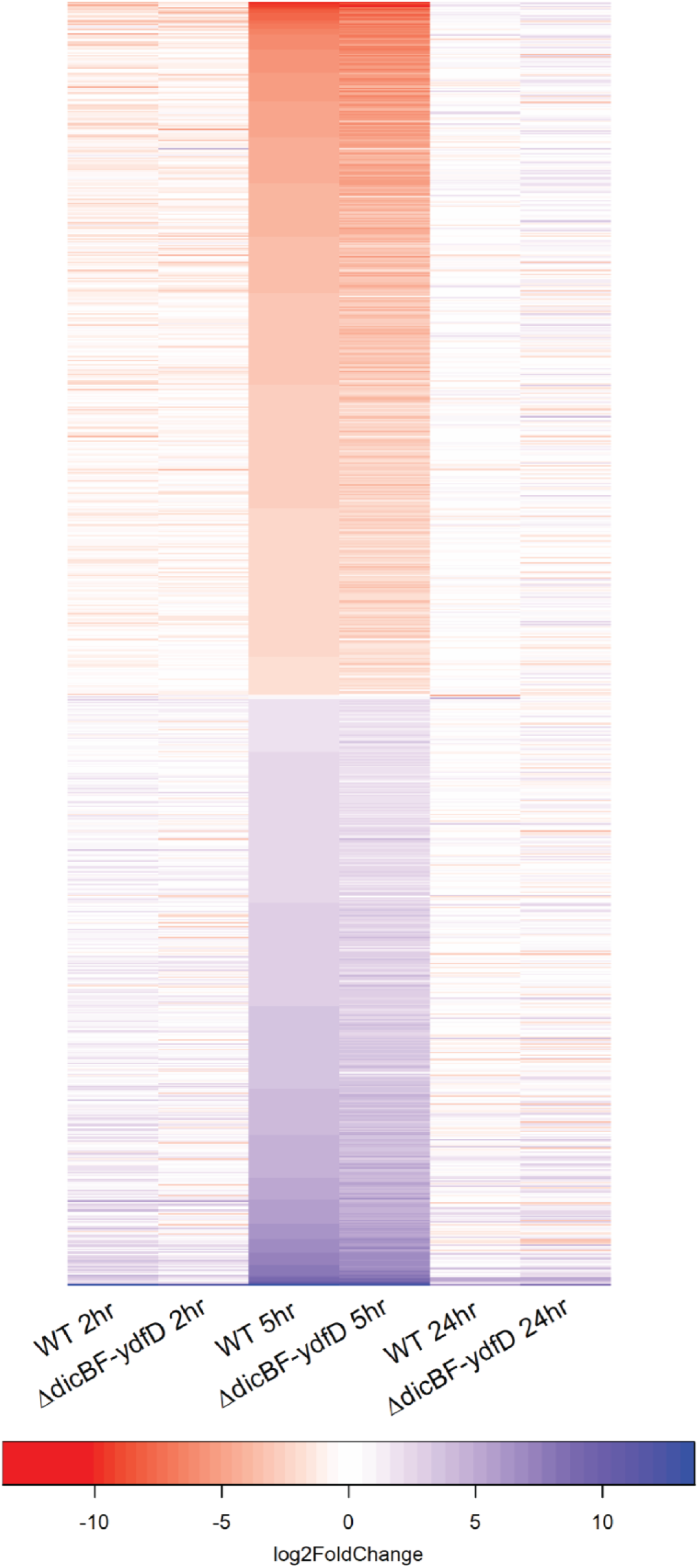
DicC-Mediated Differential Expression is Independent of *dicBF-ydfD*. Heatmap of log_2_ fold-change for genes differentially expressed under *dicC* induction compared to wild-type strain with empty plasmid induction. Genes are sorted by fold-change in wild-type background at 5-hours, up-regulation is in blue, and down-regulation is in red. Each pair of columns show fold-change of DE genes with *dicC* induction in wild-type background (left) and Δ*dicBF* background (right) across the three time points tested (2 hours: left; 5 hours: middle; 24 hours: right). In each comparison, direction of fold-change is consistent whether in the wild-type or Δ*dicBF* background.

### DicC Induction Causes a Growth Defect Independent of DicBF-YdfD and Qin

A growth defect caused by *dicC* induction has previously been reported in *E. coli* O157:H7, and was suggested to be a result of DicC inducing the Qin prophage gene *dicB* [6]; however, this effect of *dicC* induction on growth was not tested in a Δ*dicB* strain. We sought to determine whether the defect could be due to DicC regulation of host genes, regulation of *dicB*, or any other genes within the Qin prophage. We measured growth of wild-type

*E. coli* K-12 MG1655 and an isogenic *dicBF*-*ydfD* deletion strain, with and without expression of *dicC* from an inducible plasmid. The previous report showed that *dicC* induction resulted in minimal growth for 6 hours, at which point the cells begin to grow. In our experiment, both wild-type and the Δ*dicBF*-*ydfD* strains grew the same for 3 hours post-subculture, regardless of *dicC* induction. After 3 hours, both wild-type and Δ*dicBF*-*ydfD* strains showed slowed growth due to expression of *dicC* from the inducible plasmid (Figure 4A). The growth rate increased in both genetic backgrounds after ∼8 hours post-subculture until reaching stationary phase at ∼20-hours post-subculture, with the Δ*dicBF* strain growing slightly faster and growing to a slightly higher density than the corresponding wild-type strain (OD600 = 0.6 vs 0.5, respectively). The non-induced cells for both wild-type and Δ*dicBF* strains largely stopped growing at 6 hours post-subculture, reaching an OD600 of ∼0.45. These data demonstrate that the DicC-mediated growth defect is independent of *dicBF* and *ydfD* but suggest that *dicBF* and *ydfD* may impact growth recovery.

**Figure 4.**
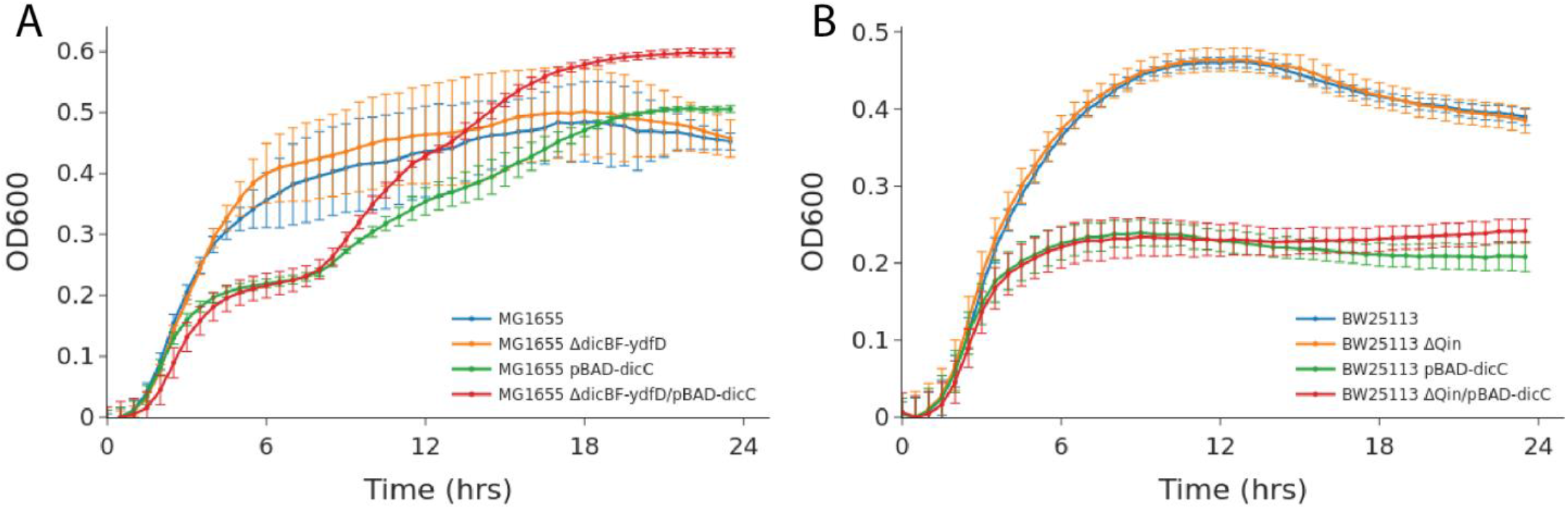
DicC-Induced Growth Defects are Independent of dicBF and Qin. Growth curves demonstrating the effect of DicC induction in various *E. coli* backgrounds. Data was collected in triplicate every 30 minutes for 24 hours; curves are plotted as the average of the replicates with error bars showing the standard deviation. **A)** The K-12 MG1655 background grows roughly the same in its WT form (blue) and when the toxic *dicB, dicF*, and *ydfD* genes are deleted (orange). Around 2-3 hours, strains with *dicC* induction begin to slow their growth. This effect is the same regardless of whether in a wild-type (green) or Δ*dicBF-ydfD* background (red) and begins to subside around 8-hours. At this point, the Δ*dicBF-ydfD* background resumes growth at a rate slightly faster than the wild-type *dicC*-induced background. **B)** The BW25113 background also grows nearly identically with (blue) or without (orange) the Qin prophage region. The key difference in this background is that the growth defect beginning around 2-3 hours is not transient, *dicC* induction impacts growth identically regardless of whether the Qin prophage is present (green) or absent (red).

To determine whether the growth defect associated with *dicC* expression could be attributed to *dicC* regulation of Qin prophage genes other than *dicBF* and *ydfD*, we measured growth of wild-type and a Qin prophage deletion mutant of *E. coli* BW25113 from [7], with and without *dicC* induction (Figure 4B). As with the MG1655 strain background, we observed slowed growth after 3 hours post-subculture in strains expressing *dicC* from an inducible plasmid; there was no difference in growth between the wild-type and prophage deletion strain backgrounds. Overall, these results show that the growth defect associated with high-level expression of *dicC* is largely independent of genes within the Qin prophage, strongly suggesting that DicC regulation of host genes is responsible for the growth defect.

## Discussion

Here, we report the ability of a cryptic prophage TF, DicC, to hijack the regulatory program of the host cell with significant phenotypic consequences. We identified hundreds of bound DNA regions for the Qin TF DicC when it was induced from a plasmid. Many of these regions could be associated with differential expression that is independent of the toxic *dicBF* genes known to inhibit cellular growth and suggested to be active under DicC induction, as well as the *ydfD* gene known to cause cell lysis. While many of these regions are bound relatively weakly compared to the most-enriched region at the *dicC* promoter, low-affinity binding has gradually gained attention over the last decade as it is capable of driving significant phenotypic changes [16-18]; thus, weak binding may still have regulatory implications. We also confirmed that the growth defect previously seen under DicC induction is independent of *dicBF* and *ydfD*, contrary to a prior report that suggested activation of *dicB* by DicC as the driver of the growth defect [6]. Further, we showed that the DicC-induced defect is independent of the Qin prophage entirely.

While some regulatory interactions for prophage TFs have been previously described [21-24], they are largely limited to genes within the prophage. To our knowledge, there has only been one other study that examined the possibility of prophage TF regulation of host genes, where the λ phage repressor *cI* was shown to impair growth of *E. coli* grown with succinate as a carbon source by directly repressing the host *pckA* gene [25]. Here we have shown that DicC is capable of broad host regulation, and our compendium of binding profiles that led to this work also identified widespread binding of other prophage TFs including DicA and YagI (Supplementary Figure 2), suggesting that this phenomenon may be common.

Analysis of expression data showed that *dicC* induction resulted in repression of *dicA*, consistent with previous reports [6, 19]. The presence of a DicC binding site at the divergent promoter between *dicC* and *dicA* also supports direct repression of *dicA* by DicC and is consistent with a previous report that these TFs appear to be wired as a bistable switch [10]. Notably, we were not able to confirm regulation of *dicB* and *dicF* through our expression analysis. These were previously reported to be induced under DicC induction [6, 10], and the presence of binding sites at the promoters known to transcribe these genes suggests they may be regulated in a condition-specific manner.

Our expression data also provide insight into previously observed *dicC* induction phenotypes, including a decrease in acid and oxidative stress resistance, β-lactam antibiotic resistance, and the growth defect caused by *dicC* induction [6]. We identified several differentially expressed host-encoded binding target genes that could explain these phenotypes. First, the decrease in acid resistance is consistent with the down-regulation of Glutamate-Dependent Acid Resistance genes, as GDAR is the main acid resistance system in *E. coli* [26]. Notably, *gadE*, the gene encoding the primary TF responsible for GDAR regulation, was 75-fold repressed at 5-hours post-induction. We also found the primary decarboxylases that facilitate GDAR by converting glutamate and a hydrogen to 4-aminobutanoate and carbon dioxide, *gadA* and *gadB*, to be 617- and 699-fold repressed, respectively, at 5-hours post-induction. The glutaminase that converts glutamine into glutamate, *glsA* was 405-fold down-regulated at 5-hours post-induction; and finally, *gadC*, the transporter responsible for importing glutamine and glutamate, which are critical for GDAR, and exporting 4-aminobutanoate, was 898-fold repressed at 5-hours post-induction. Each of these genes were slightly repressed at 2-hours post-induction as well (from 2.7-fold to 19.4-fold down-regulated), but these changes were not statistically significant after multiple test correction.

The decrease in oxidative stress resistance with *dicC* induction is consistent with the down-regulation of peroxidase *tpx* (11-fold and 174-fold repressed at 2- and 5-hours post-induction, respectively), as a *tpx* deletion was previously shown to be more sensitive to this stress [27].

The reduction in β-lactam antibiotic stress resistance from *dicC* induction is consistent with the observed up-regulation of outer membrane porin *ompF* (5-fold up-regulation at 5-hours post-induction). This gene facilitates entry of β-lactams into the cell, and deletions of *ompF* have been shown to increase antibiotic resistance [28]; thus up-regulation of this gene is consistent with increased susceptibility to β-lactams.

The growth defect resulting from *dicC* induction is consistent with the observed up-regulation of *ydeP* (20- and 47-fold up-regulated at 2- and 5-hours post-induction, respectively), as *ydeP* induction has been shown to decrease growth rate in exponentially growing cells [29]. Curiously, a *ydeP* deletion was also found to be more sensitive to acid stress [30], however the mechanism by which *ydeP* might confer acid resistance is not known and it is unclear if *ydeP* up-regulation could offset acid susceptibility due to GDAR repression. A second gene, *symE*, that was 12-fold up-regulated at 5-hours post-induction has been shown drive nucleoid condensation, which can damage DNA and block replication, as well as disrupt RNA synthesis [31], ultimately hindering cell growth.

The measurement of the growth defect caused by *dicC* induction allowed us to confirm that it was both independent of the *dicB, dicF* and *ydfD* genes (where *dicB* was previously suggested to cause the growth defect under *dicC* induction [6]), as well as the Qin prophage more broadly. Our data also suggest that some combination of *dicB, dicF*, and *ydfD* impact growth recovery after *dicC* induction. However, there was no difference in growth recovery between wild-type BW25113 and the Qin prophage deletion, suggesting that the impact of *dicB, dicF*, and *ydfD* on growth recovery is indirect.

We observed the effect of *dicC* expression from an inducible plasmid rather than endogenous expression. Nonetheless, analysis of public RNA-Seq datasets found *dicC* to be highly expressed when cells were exposed to norfloxacin or cold shock (data not shown), suggesting potential inducers. One study has implicated another Qin gene, *rem*, as a potential mechanism for alleviating *dicA* repression [10], which could enable *dicC* expression, though *rem’s* inducer is unknown.

Overall, our results speak to a closer relationship between TFs encoded by cryptic prophages and their host cells than previously thought. The ChIP-Seq data from our previous work identified hundreds of binding sites for 3 CP-encoded TFs [15]. Together with the results in this manuscript, these data suggest that prophage-encoded transcription factors have the potential for inducing phenotypic changes through regulation of host-encoded genes. Phenotypes previously associated with *E. coli* prophages have mostly been understood locally through the effects of prophage-encoded genes, but our results suggest that prophage-encoded TFs may allow prophages to impact physiology more broadly.

## Methods

### Strains and Plasmids

A list of strains and plasmids is available in Supplementary Material.

#### Inducible Tagging for in vivo ChIP-Seq

Inducible pBAD24 *dicC* constructs were generated according to previous methods [15]. Briefly, *dicC* was PCR-amplified from *E. coli* genomic DNA using primers that target *dicC* (except for the start and stop codons) and contain overhangs with identity to the pBAD insertion site. Plasmid was linearized with *KpnI* (NEB #R0142) and purified with a PCR Purification Kit (Qiagen #28104). PCR fragments and linearized vector were ligated with the HiFi Assembly Mix (NEB #E2621) and ligation was transformed by heat shock into *E. coli K-12* MG1655.

#### Creation of Deletion Strains

The *dicF, dicB*, and *ydfD* three-gene locus was replaced by *thyA* using the FRUIT recombineering method [32] to generate the Δ*dicBF*-*ydfD*::*thyA* strain.

### ChIP-Seq

ChIP-Seq was carried out according to our paper in submission [15]. Briefly, the *E. coli* K-12 MG1655 pBAD *dicC* strain was grown in 5 mL M9 minimal media with 0.4% glycerol as a carbon source and ampicillin selection in a vented tube at 30°C with 250 rpm shaking overnight. The following day, cells were subcultured 1:100 in 50 mL fresh media and allowed to grow to log-phase, at which point arabinose was added to a final concentration of 0.2% for DicC induction. Cells continued to grow for 1 hour before formaldehyde was added to a final concentration of 1%, and flasks were shaken at 100 rpm for 30 minutes at RT to crosslink TF-DNA interactions. Crosslinking was ceased with the addition of glycine to a final concentration of 250 mM and incubating for 15 minutes longer at RT with 100 rpm shaking. Crosslinked cells were washed twice with 50 mL 1X PBS. After the second wash, cells were resuspended in a mixture of 600 µL Buffer 1 (20 mM HEPES Potassium Salt pH 7.9, 50 mM KCl, 0.5 mM DTT, 10% glycerol) + Protease Inhibitor (Sigma Aldrich #04693116001), transferred to Covaris-compatible tubes, and left on ice prior to cell lysing and DNA shearing via sonication. This was achieved using the Covaris S2 sonication platform for 12 minutes with amplitude 20%, intensity = 5, and 200 cycles/burst. Sonicated lysate was transferred to a sterile 1.5 mL tube and pelleted by centrifugation at 13,000g for 10 minutes in a 4°C centrifuge. Supernatant was transferred to fresh 1.5 mL tubes and salt concentration was adjusted by adding Tris-HCl pH8, NaCl, and NP-40 to final concentrations of 10 mM, 150 mM, and 0.1%, respectively. Next, we added 5 µL of Anti-FLAG antibody (Sigma-Aldrich #F1804) and left on a rocking platform at 4°C overnight. The following day, immunoprecipitation was carried out by first washing 50 µL Protein G Agarose beads (Thermo Scientific #20399) with 1 mL IPP150 buffer (10 mM Tris-Hcl pH8, 150 mM NaCl, 0.1% NP-40). Beads were spun in a centrifuge at 2,000g for 2 minutes at RT and supernatant was discarded. Antibody-containing lysate from the previous day was then added to the beads, and tubes were returned to the rocking platform at 4°C for 30 minutes, before being moved to RT for another 90 minutes of rocking. Beads were washed 5 times with IPP150 buffer, followed by 2 washes with 1X TE buffer. After the last wash, 150 µL EBB buffer (50 mM Tris-HCl pH 8, 10 mM EDTA, 1% SDS) was added to the bead pellet and tubes were incubated at 65°C for 15 minutes, prior to pelleting at 2,000g for 5 minutes at RT. Supernatant from the first elution was transferred to a sterile 1.5 mL tube, and a second elution was carried out by adding 100 µL 1X TE + 1% SDS, incubating for 5 minutes at 65°C, and pelleting again. The second elution was pooled with the first, and 12.5 µL Proteinase K (Sigma-Aldrich #P4850) was added. Tubes were incubated at 37°C for 1 hour to digest DNA-bound proteins, followed by incubation at 65°C overnight to deactivate the proteinase. The following day, ChIP DNA was purified using Qiagen’s PCR Purification Kit (Qiagen #28104). Sequencing libraries were generated from ChIP-DNA using the NEBNext Ultra Library Prep Kit for Illumina (NEB #E7645) and NEBNext Multiplex Oligos for Illumina dual-index primers (one of #E6440, #E6442, #E6444, #E6446, #E6448). Concentrations and size distributions of prepared libraries were analyzed with the Agilent Bioanalyzer 2100 using High-Sensitivity DNA chips. Libraries were normalized to 4 nM and pooled for sequencing, in cases where libraries were less than 4 nM, an equimolar volume was added to the pool.

### ChIP-Seq Analysis

Analysis of ChIP-Seq data was carried out according to our prior method [15]. Briefly, we performed quality control on raw reads with FastQC [33] prior to pruning Illumina adapter sequences with Cutadapt [34] and aligning to *E. coli* (Genbank Accession U00096.3) using Bowtie2 [35]. We calculated per-base coverage by determining the number of reads aligning to each strand at each position. Coverage values were used to identify statistically enriched regions of coverage using SPAT [36]. We applied several filters to these regions, including: (1) filtering for the expected signature of forward and reverse coverage; (2) filtering for background enrichment in control experiments; and (3) filtering for ChIP-Seq-specific artifacts that are not apparent in controls. After filtering likely spurious regions, we performed ChIP-Seq-specific quality control to verify tagging of the correct TF. The final set of enriched regions reported here for *dicC* were generated by combining any overlapping regions across four replicates. Regions that were identified in more than one replicate were included in our set.

### Assignment of Potential DicC Targets

To determine which genes might be regulated by a DicC binding site, we obtained a list of annotated transcription units (TUs) from RegulonDB [12]. For each position of the genome, we identified the closest TU on either side of the coordinate within 400 bp, such that at most two TUs could be regulated from a given position. In cases where more than one TU had the same strand and start position, we used the longest TU. For each DicC binding region, we identified the approximate center of the binding site as the position with maximum enrichment and assigned potential regulatory targets based on the TUs assigned to that position. Each gene inside of the assigned TU was considered a putative direct target of DicC.

### RNA-Seq

RNA-Seq was carried out in duplicate following established methods [26] using the same strains described above. Cells were inoculated from glycerol stocks at 37°C in 5 mL LB (and ampicillin when pBAD was present) with 250rpm shaking. The following day, cells were subcultured into 5 mL fresh LB + 0.2% arabinose for induction Total RNA was extracted from cells after 2-, 5-, and 24-hours of growth using Qiagen’s RNeasy Mini Spin Kit (Qiagen #74104) and subjected to DNAse digestion by combining 5 µg RNA with 1 µL TURBO DNAse enzyme, 5 µL 10X Reaction Buffer (Invitrogen #AM2238), and raising the total volume to 50 µL with sterile water and incubating at 37°C for 1 hour. DNAse-digested RNA was cleaned up using RNAclean XP beads (Beckman Coulter #A63987) by adding 90 µL beads (for a 1.8X bead:sample ratio) and incubating for 10 minutes. Samples were moved to a magnetic rack for 5 minutes and supernatant was discarded. Beads were washed twice on the magnetic rack with 200 µL 70% ethanol. After the second wash, beads were allowed to dry on the magnetic rack for up to 10 minutes before RNA targets were eluted in 27 µL RNAse free water. RNA concentrations were determined with Qubit RNA HS Assay Kit (Thermo Fisher #Q32855). Extracted RNA was converted to NGS libraries using Zymo-Seq RiboFree Total RNA Library Kit (Zymo #R3003) using 1 µg RNA as input. Quality control and library molarities were assessed with Agilent’s Bioanalyzer 2100, and libraries were pooled in the same way used for ChIP-Seq samples. Sequencing was performed on Illumina’s NextSeq 2000 platform with 75 bp single reads.

### RNA-Seq Analysis

#### Sample Pipeline

RNA-Seq analysis was performed following previously published methods [37]. Briefly, we started with the same steps of raw read quality control with FastQC and adapter trimming with Cutadapt that were used in the ChIP-Seq pipeline above. We used HISAT2 [38] for alignment of RNA-Seq reads with the “—rna-strandedness” parameter set to “RF”, since reads produced by the Zymo kit used are expected to align to the opposite strand of the gene. Transcripts were first quantified using HTSeq [39] to determine the number of read counts aligning to each gene, then additionally quantified with Cufflinks [40] to obtain FPKM and TPM measurements of gene expression.

#### Differential Expression

Differential expression of genes was assessed using DESeq2 [41]. Briefly, raw counts from each sample were gathered into an expression matrix. Genes with at least 10 counts in at least 2 samples were retained, and expression values across samples were normalized with the DESeq “median of ratios” method. Contrasts were configured to compare the effect of DicC induction vs empty plasmid at each time point, or the effect of *dicBF* knockout with DicC induction vs empty plasmid at each time point. Finally, for each comparison, we defined differentially expressed genes as those with an adjusted pvalue of 0.05 or less (using default Benjamini-Hochberg correction), and an absolute log_2_ fold change of at least 2 (4-fold up-or down-regulation).

### Growth Curves

Growth curves were measured to analyze the *dicC*-induced growth defect using the strains described above. Cells were initially inoculated in 5 mL LB (with ampicillin selection where necessary) and grown overnight aerobically at 37°C with 250 rpm shaking. The following day, a 96-well plate was prepared for growth curves by adding 198 µL of LB (along with 0.2% arabinose and/or ampicillin where necessary) to each well, and 2 µL of cells from the overnight cultures. OD600 measurements were taken every 30 minutes for 24 hours using a TECAN Infinite M200 Pro heated to 37°C with shaking in between readings. Each experimental group was measured in triplicate and the average of three replicates at each time point were reported with standard deviation as error bars.

## Supporting information

Enriched GO Categories

Differential Expression Values

Enriched GO Categories (with genes, in .json format)

## Acknowledgements

We acknowledge Boston University’s Microarray and Sequencing Resource Core Facility which performed NGS for ChIP-Seq and most RNA-Seq experiments, as well as the DAMP lab, which performed NGS for some of the RNA-Seq experiments. We acknowledge the Wadsworth Center Media and Glassware Facility for providing supplies to perform strain construction. We acknowledge BioRender which assisted in the generation of Figure 1A.

## Funding

National Institutes of Health grant 5R01GM131643-04 (PL, VHT)

National Institutes of Health grant 5R01GM114812-04 (PL)

National Institutes of Health grant 5R01EB029795-02 (PL, VHT)

Maximizing Investigators Research Award 5R35GM144328 (JTW)

Universidad Nacional Autónoma de México (LGR, VHT, JCV)

## Authors Contributions

Conceptualization – PL, JEG, JCV, JTW

Methodology – PL

Software – PL

Validation – PL, VHT

Formal Analysis – PL

Investigation – PL, VHT

Resources – PL, VHT, LGR, AS

Data Curation – PL, LGR

Writing – original draft preparation – PL

Writing – review and editing – PL, VHT, LGR, JCV, JTW, JEG

Visualization – PL

Supervision – JCV, JTW, JEG

Project Administration – PL, JEG

Funding Acquisition – JCV, JTW, JEG

## Competing Interests

JEG is a co-founder of Biosens8, Inc.

## Data and Materials Availability

Raw data will be made available in NCBI’s Gene Expression Omnibus (GEO).

Processed data will be uploaded to RegulonDB.

