## Supplementary Material for "A Cryptic Prophage Transcription Factor Drives Phenotypic Changes via Host Gene Regulation"

### Supplementary Figures

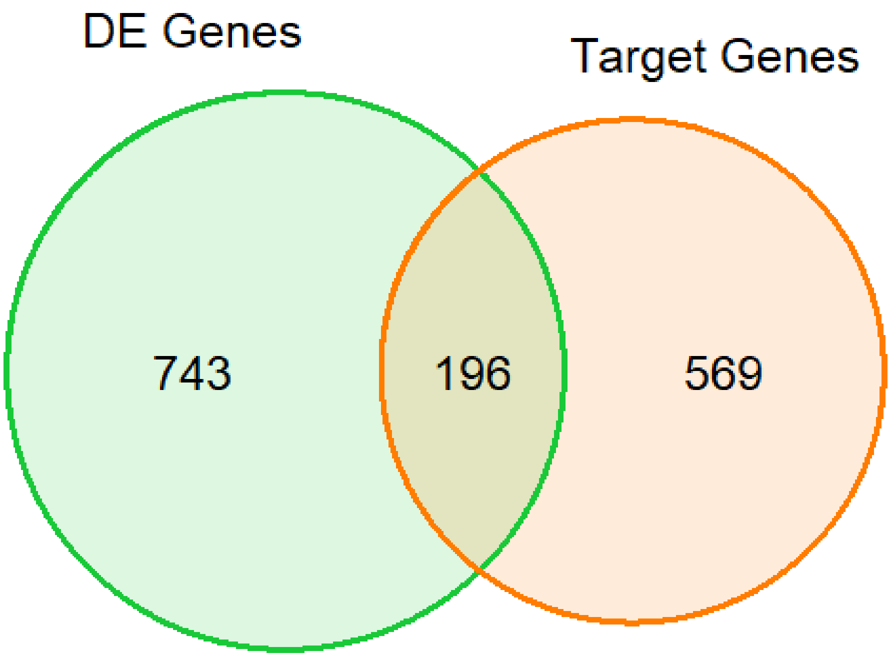


**Figure S1 Venn Diagram of DE and target genes.** Venn diagram displaying the overlap between differentially expressed genes (939 total) and putative direct *dicC* targets that were detectable in expression analysis (765 total). There were a total of 196 direct targets that were also DE.

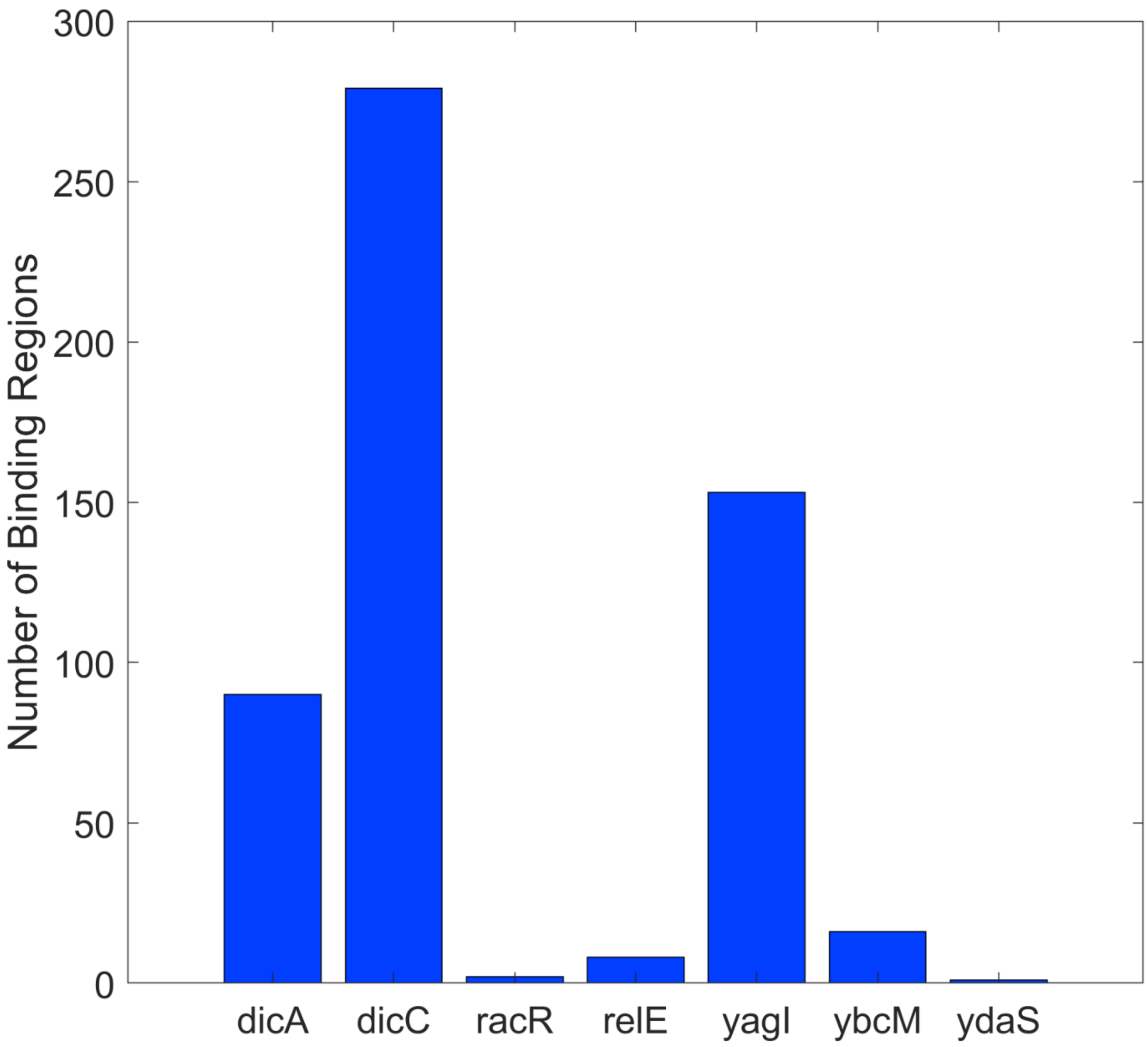


**Figure S2 Other Prophage TFs Have Potentially Wide Regulons.** Bar chart for 7 prophage TFs (including *dicC*) that were studied in our recent work [1]. Two other prophage TFs, *dicA* and *yagI* were found to bind many regions across the genome, though to a lesser extent than *dicC*. This suggests that other prophage TFs may also be capable of broad regulation of the *E. coli* host.

### Supplementary Tables

| Strain | Plasmid | Source |
|---|---|---|
| MG1655 | - | Wade Lab |
| MG1655 pBAD | pBAD24-FLAG0 | Wade Lab |
| MG1655 pBAD-dicC | pBAD24-dicC-FLAG0 | Wade Lab |
| MG1655 ΔdicBF-ydfD | - | Wade Lab |
| MG1655 ΔdicBF-ydfD/pBAD-dicC | pBAD24-dicC-FLAG0 | Galagan Lab |
| BW25113 | - | Microbiologics NCTC 14364 |
| BW25113 ΔQin | - | Microbiologics NCTC 14370 |
| BW25113 pBAD-dicC | pBAD24-dicC-FLAG0 | Galagan Lab |
| BW25113 ΔQin/pBAD-dicC | pBAD24-dicC-FLAG0 | Galagan Lab |

**Table S1 Strains and Plasmids**
